# Causal network perturbations for instance-specific analysis of single cell and disease samples

**DOI:** 10.1101/637710

**Authors:** Kristina L. Buschur, Maria Chikina, Panayiotis V. Benos

## Abstract

Complex diseases involve perturbation in multiple pathways and a major challenge in clinical genomics is characterizing pathway perturbations in individual samples. This can lead to patient-specific identification of the underlying mechanism of disease thereby improving diagnosis and personalizing treatment. Existing methods rely on external databases to quantify pathway activity scores. This ignores the data dependencies and that pathways are incomplete or condition-specific.

ssNPA is a new approach for subtyping samples based on *deregulation* of their gene networks. ssNPA learns a causal graph directly from control data. Sample-specific network neighborhood deregulation is quantified via the error incurred in predicting the expression of each gene from its Markov blanket. We evaluate the performance of ssNPA on liver development single-cell RNAseq data, where the correct cell timing is recovered. In all analyses ssNPA consistently outperforms alternative methods, highlighting the advantage of network-based approaches.

## Introduction

Gene expression profiling by RNA-sequencing has become routine tool in biomedical research. Similarly, on the clinical side, RNA-seq has now been introduced as a cost-effective diagnostic tool ^1,2^. Moreover, recent technological advances have made the assessment of gene expression at single cell level (scRNA-seq) feasible, opening new avenues to developmental biology and the study of dynamic networks ^3–5^. Consequently, the number of large RNAseq datasets keeps growing with hundreds or thousands of samples representing a single clinical or cellular condition. As a result, the scientific questions have shifted away from simple differential expression to characterizing the molecular heterogeneity of disease phenotypes. One simple way to characterize sample heterogeneity is via clustering and/or dimensionality reduction. This approach will often reveal distinct sample groups within the population but it treats all genes equivalently, ignoring the fact that genes are organized in regulatory networks. On the other end of the spectrum there has been considerable development in methods that quantify pathway activation on a single sample level (ssGSEA ^6^, PLAGE ^7^, GSVA ^8^, Pathifier ^9^). However, these methods rely heavily on existing pathway information (e.g., from KEGG, BioCarta, The Nature Pathway Interaction Database), which may be incomplete, not well annotated or irrelevant to the studied phenotype or condition. Other methods (e.g., ^10^) quantify a sample-to-sample similarity with the aim of identifying similarities and differences between cell functions.

In this paper, we present a different approach for assessing, qualitatively and quantitatively, how the gene network from a set of control samples is perturbed in a newly presented single sample. Our approach, <u>Single Sample Network Perturbation Assessment</u> (ssNPA), uses causal modelling to first learn the gene expression interaction network from a set of reference samples. For each new sample the method then assesses which parts of the “reference sample network” are deregulated.

Causal graphs have been used in the past to learn gene networks from expression data ^11–13^ or gene features that are highly predictive of certain phenotypes ^14–18^. Our ssNPA approach learns a causal graph from expression data and for every gene it builds a predictive model based on its Markov blanket. Applying the model to a new sample produces a vector of residuals which quantifies the network level gene dysregulation (NLGD). The NLGD vectors can then be used to cluster samples into groups and assess their group characteristics (e.g. developmental time, survival, molecular mechanisms of phenotype, etc.) or to assign an individual patient to a disease subcluster. We use this property to evaluate ssNPA on existing datasets, for which the ground truth is known. Specifically, we show that ssNPA separates well the developmental trajectory and the differentiated cell type in a mouse liver cell development scRNA-seq dataset ^19^, demonstrating that ssNPA can separate the samples in the corresponding datasets according to patient survival and molecular subtypes with better accuracy than alternative approaches.

## Results

### ssNPA Algorithm Description

ssNPA learns the global gene expression network as a directed (causal) graph from a set of reference samples using FGES ^20^. FGES calculates a directed acyclic graph (DAG) over all data by maximizing the Bayesian Information Criterium (BIC) score of the data given the model (network). The BIC score is given by the formula:

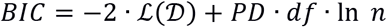

where 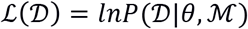 is the maximum log-likelihood of the data given the model and its parameters; *PD* is a penalty value (“penalty discount”) that controls sparsity (*PD*=1 in the standard BIC definition); *df* is the degrees of freedom; and *n* is the sample size. This score is decomposable and the total BIC of the graph is the sum of the BIC of its nodes and their parents. FGES starts with an empty graph then adds single edges while the BIC score increases. Next, the algorithm removes single edges while the BIC score increases.

The Markov blanket of a gene *G_i_* (MB(*G_i_*)) consists of the parents, children and spouses of *G_i_* in the graph. Once the graph has been learned from the reference (control) samples, then ssNPA uses the Markov blanket around each gene, *G_i_*, to build a predictor of its expression. This is because in a directed graph:

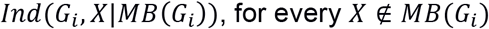

Therefore, a highly predictive regression model can be learned for each gene:

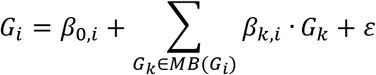

Then for each new sample this model can be used to calculate the deviation of the expression of *G_i_* in this sample compared to the reference samples. So, the new sample can be represented as a vector of deviations of expression of every gene from the reference samples. Given that genes are connected through the network of interactions, in this way, we assess both the topology and the magnitude of network perturbations. This procedure is summarized in **Figure 1**.

**Figure 1.**
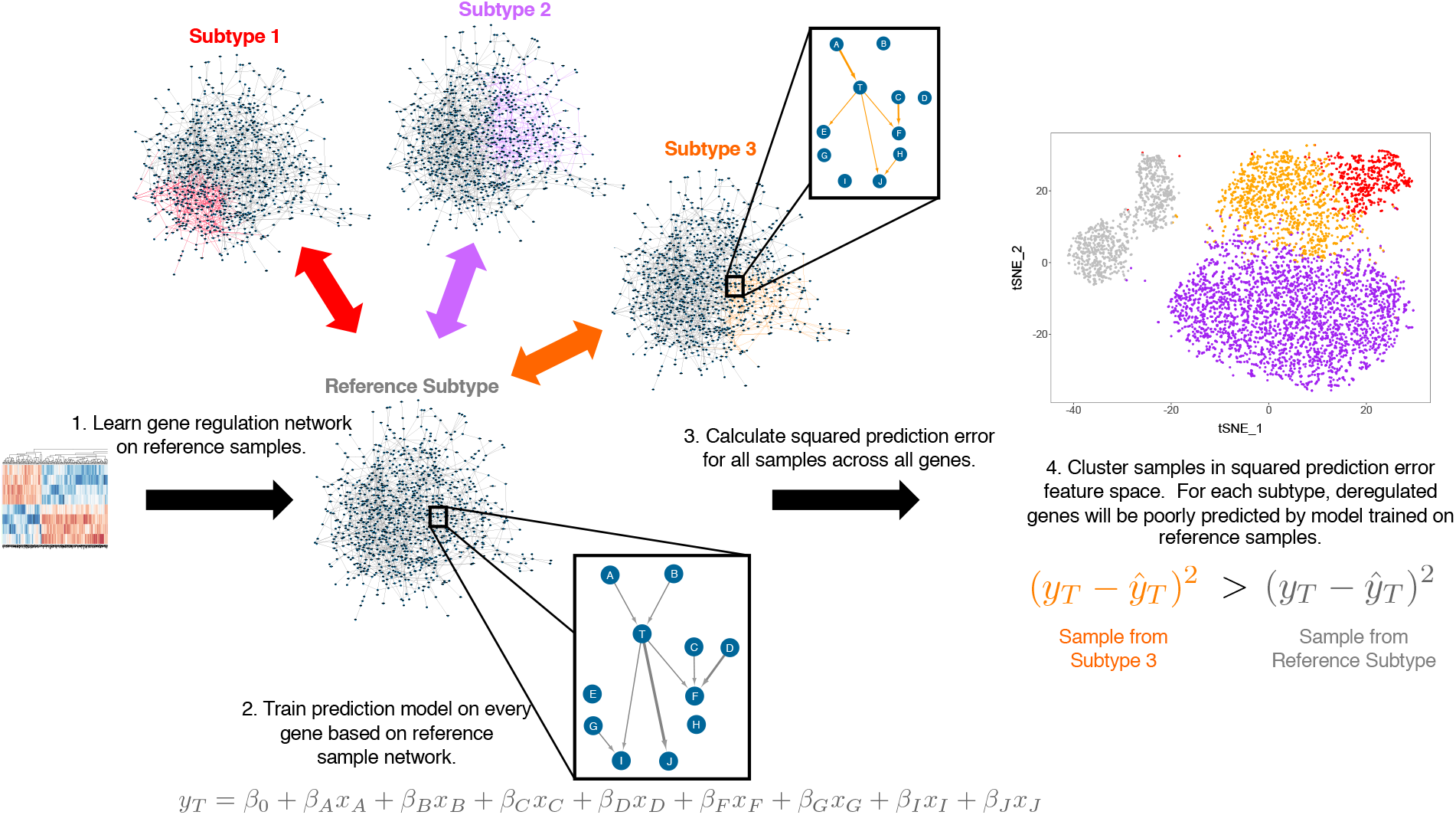
Overview of the single-sample network perturbation assessment through causal network (ssNPA) algorithm.

For comparison purposes, we also implemented ssNPA-LR, in which causal learning is substituted by lasso regression, resulting in an undirected graph. The ssNPA analysis procedure has the following steps:

1. *Data preparation*. For speed and accuracy, in this paper we selected the top 3,000 most variant genes for scRNA-seq or RNA-seq data. The RNA-seq counts were transformed to log2 counts per million through mean-variance modeling by the voom function (Limma v. 3.32.10) ^21^.
2. *Reference samples*. For disease data, we used the controls as reference sample set. For the liver scRNA-seq, we tested each stage (as determined by external cell markers) as potential reference group.
3. *Gene network learning (ssNPA)*. A directed graph was learned on the expression data for the reference group of samples (FGES algorithm) ^20^. For this work, we scan over a number of PD values in the range [4, 12] and we choose a PD for each dataset that balances grouping the reference samples together while not overfitting. The Markov blanket around every gene in this network is used for predicting its expression on any given sample; and deviation from the observed value is a measure of network perturbation.
4. *Feature selection (ssNPA-LR)*. In this case, we used the glmnet package in R (v. 2.0.16) to learn a lasso regression prediction model for every gene across the reference samples ^22^. We chose each sparsity parameter (λ) with 10-fold cross validation, selecting the value of λ corresponding to minimum mean cross-validated error.

### ssNPA correctly identifies embryonic stage and cell type in murine liver cells from single cell RNA-seq data

We used a recently published liver development scRNA-seq dataset to test ssNPA and compare it to other methods. This dataset is composed of multiple types of liver cells samples at a series of developmental timepoints. The early hepatoblast cell differentiates into two lineages (hepatoblasts and cholangiocytes). In this dataset the time point and cell-identity is experimentally controlled and thus can serve as the ground truth. We hypothesize that information regarding the cell-type and developmental stage is reflected in the gene expression data.

When gene expression data used directly for clustering ^23^ we identified six clusters (Supplementary Figure S1A), which separated well the extreme developmental time points: cells measured at day E10.5 and hepatocytes from day E17.5 (**Figure 2A**). However, all of the differentiated cholangiocytes were grouped together in a single cluster and although they were somewhat stratified within the cluster, their embryonic stage was not distinguishable. The remaining clusters contained a mix of intermediate timepoints (days) and hence they did not accurately represent the developmental trajectory.

**Figure 2.**
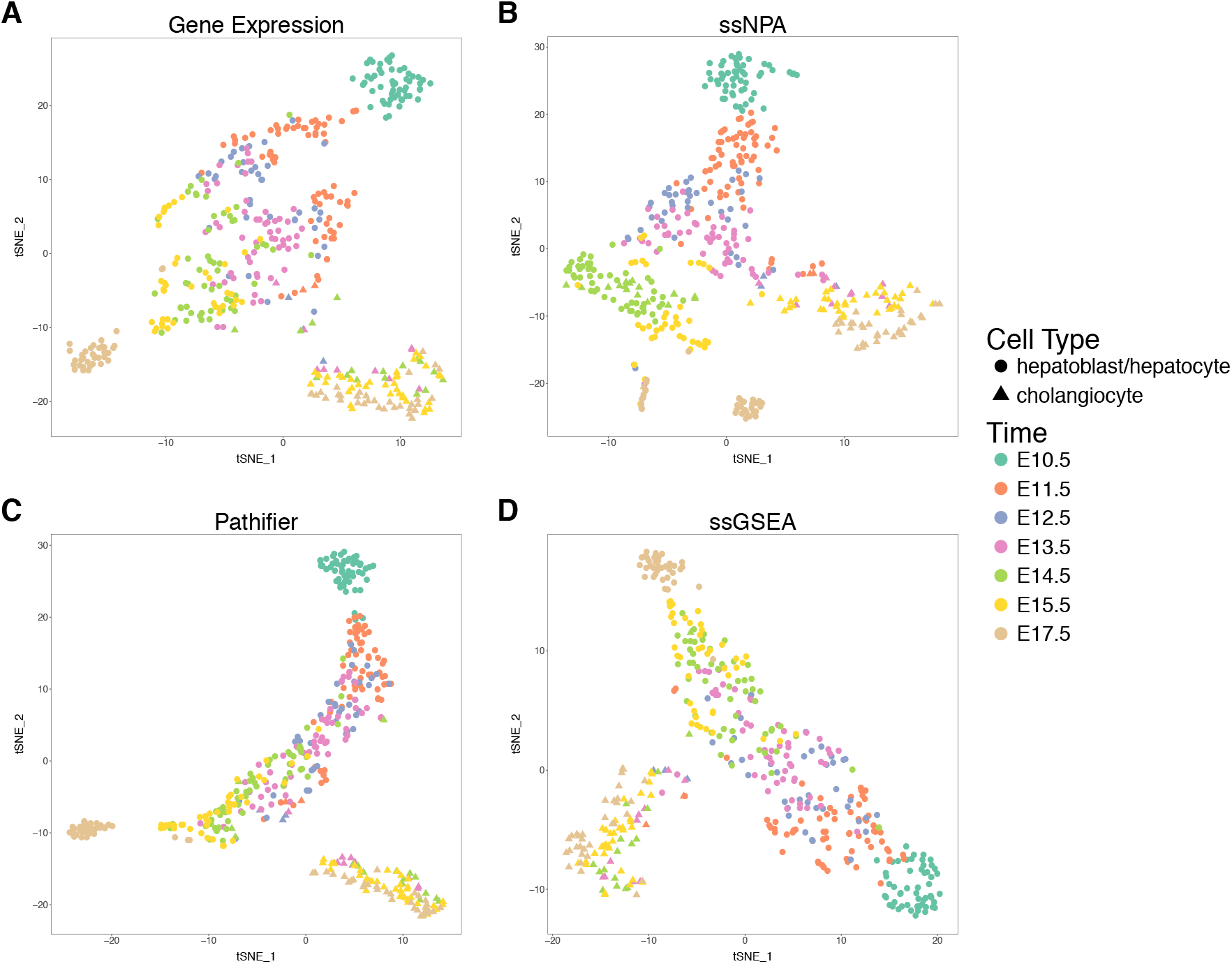
Comparison of how well (A) gene expression, (B) ssNPA, (C) Pathifier, and (D) ssGSEA separate murine liver cell scRNA-seq samples by developmental stage and cell type. ssNPA was used with the E14.5 cells as the reference set and PD=5. Pathifier was also applied with the E14.5 cells as the reference set. Clustering for every method was performed with the first ten principal components.

By contrast, the six identified clusters based on the network perturbation features of ssNPA (Supplementary Figure S1B) separated well all stages (**Figure 2B**). In particular, hepatoblasts from days E10.5 and E11.5 as well as mature cholangiocytes and E17.5 hepatocytes were separated into four distinct clusters. Since there was not an obvious reference set of samples in this dataset, we examined the utility of each group as a potential reference. We additionally evaluated a range of PD (penalty discount) parameter values for FGES [4,12]. We found the late intermediate stages (E14.5 or E15.5) to show better performance than the extremes, when they were used as reference set, while the choice of PD had less impact on clustering performance (Supplementary Figure S2).

Next, we compared ssNPA to Pathifier and ssGSEA. Both methods quantify gene interactions, but require pathway information from an external database. For Pathifier we used the KEGG pathway database ^24^. It also identified six distinct clusters (Supplementary Figure S1C), but with the exception of E10.5, it did not separate the developmental stages very well (**Figure 2C**). All of the intermediate stage hepatocytes were mixed together and distributed in three clusters. Furthermore, the runtime of Pathifier was very long compared to ssNPA (on the order of hours compared to minutes), Finally, we tested ssGSEA which calculates a gene set enrichment score for every sample. We used ssGSEA with default parameters and the default gene sets from the C2 collection of the Molecular Signatures Database version 3.0 ^25^. ssGSEA does not require the user to provide a reference set. Clustering with the ssGSEA produced five clusters (Supplementary Figure S1D), but in general, these were not well separated according to developmental time point (**Figure 2D**). The cholangiocytes were grouped into one cluster but the largest cluster contained the hepatoblasts from E10.5 and E11.5. The hepatocytes from E17.5 were grouped together well in another, but the remaining hepatoblasts/hepatocytes spanning E12.5-E15.5 were mixed together and divided between two clusters.

We additionally developed and tested a variation of ssNPA, the ssNPA-LR algorithm, which uses lasso regression instead of causal learning to choose the features predicting the expression of a gene (Supplementary Figure S3A). We found five clusters (Supplementary Figure S3B), which separated well the early and late developmental stages, but most of the intermediate stage hepatocytes were grouped together in one cluster. The exception to this is the group of E14.5 hepatocyte and cholangiocyte cells, which are grouped tightly together at a large distance from the rest of the developmental trajectory. This suggests that the lasso regression approach might be overfitting to the reference group of cells despite the fact that its optimum parameter was selected through cross-validation to minimize mean error.

To quantitatively compare the clustering performances of all methods we used the normalized mutual information (NMI) and the adjusted Rand index (ARI) (**Table 1**). We found that ssNPA and ssNPA-LR clearly outperform Pathifier, ssGSEA, and gene expression (with either the top 3,000 most highly variant genes or all genes) by maximizing NMI (0.693 and 0.687, respectively). ssNPA also returned the highest ARI of these methods (0.5). However, we note a strong advantage to ssNPA over ssNPA-LR when we consider how many genes they utilized. On average, ssNPA used only 2.5 predictors for every gene, while ssNPA-LR needed 23.1 genes. Thus, we see that using directed graphs to jointly model the expression of all 3,000 genes offers a clear advantage.

**Table 1.** Comparison of different feature calculation methods. Clustering for every method was performed with the first ten principal components. E14.5 were used as reference cells for Pathifier, ssNPA and ssNPA-LR. PD=5 for ssNPA and ssNPA-LR. *MI*: mutual information; *ARI*: adjusted Rand Index.

| Method | Normalized MI | ARI | Avg number of features |
| --- | --- | --- | --- |
| **ssNPA** | **0.693** | **0.500** | **2.5** |
| Gene expr. (all genes) | 0.571 | 0.367 | NA |
| Gene expr. (top 3,000) | 0.565 | 0.369 | NA |
| Pathifier | 0.585 | 0.399 | NA |
| ssGSEA | 0.540 | 0.343 | NA |
| ssNPA-LR | 0.687 | 0.427 | 23.1 |

## Discussion

We presented ssNPA, a new method to assess gene network perturbations in each sample. The method first infers the global network from a set of reference samples using causal graph learning. In the following step given a new sample the method calculates its deviation from the reference network at every gene, thus providing information about both the topology and the magnitude of network perturbations. The perturbation feature vector can been used to cluster samples into cell or disease subtypes. We demonstrated the performance of ssNPA by using it to evaluate cluster memberships of datasets with known ground truth, specifically in liver development cells (time course scRNA-seq data). We showed that ssNPA performs better than currently used methods and from simple gene-based clustering on finding the true developmental stage and type of the cell. This showed that network perturbation features can recapitulate the time course data. In this dataset, we found that using one of the middle developmental stages (which are equadistant from both progenitor and fully differentiated extremes) as reference point allows for better results

We compared ssNPA to ssGSEA and Pathfinder, two known methods for single sample analysis. In all cases ssNPA performed better than these methods as it is evidenced by the greater agreement of the ssNPA-identified clusters to the ground truth. Using network deregulation features also differences in gene expression and also captures differences in the topology of the network of each sample from the reference samples. The better performance of ssNPA versus ssGSEA and Pathfinder might reflect the fact that the latter depend on prior knowledge that might not be very accurate or might not reflect the particular conditions in the studied dataset.

In summary, ssNPA is a new method for characterizing single samples of gene expression and offers significant advantages over existing methods. Unlike ssGSEA and Pathifier, it does not require prior pathway knowledge; it is substantially faster than Pathifier; and can be used to produce high quality sample clusters that reflect the underlying mechanisms of the disease condition or phenotype. In the future, ssNPA can be used for analyzing disease data to identify disease subphenotypes and develop personalized intervention strategies.

## Materials and Methods

### Liver Cell Development Data

A murine liver cell development scRNA-seq dataset was obtained from ^19^ (GEO:GSE90047). The experiment measured the gene expression of 447 cells over the course of embryonic days E10.5-E17.5. Cells were first sorted with fluorescence-activated cell sorting (FACS) according to the cell surface markers Delta-like (DLK) to identify hepatocytes and epithelial cell adhesion molecule (EpCAM) to distinguish cholangiocytes.

### Sample clustering

In order to better evaluate the efficiency of the various methods for single sample subtyping, we performed sample clustering using Seurat ^26^ and we examined various external characteristics of the clusters. Samples were clustered in their feature space. First, the samples are projected into principal component space. The number of principal components to retain in the projection is determined heuristically by identifying the elbow of the scree plot. Then clustering is performed with a graph-based clustering that constructs the shared nearest neighbour graph and then optimizes the modularity function ^23^. Finally, the clusters are visualized with a nonlinear dimensionality reduction (t-SNE) ^27^.

### Comparison to other methods

ssNPA methods were compared to Pathifier ^9^ and single-sample gene set enrichment analysis (ssGSEA) ^6^. All methods were tested on the same input data and reference sample selections. For Pathifier, we provided gene lists for all KEGG pathways and used the R implementation with the quantify_pathways_deregulation() function and default parameters. For ssGSEA we used the gene sets from the C2 collection of the Molecular Signatures Database version 3.0 ^25^ provided in the GSVAdata R package and the implementation of ssGSEA provided within the GSVA() function of the GSVA R package with default parameters. For fairness, we use an equal number of principal components for clustering with each method. The number of principal components is set to the maximum number of principal components identified by the elbow of the scree plot for any single method.

## DECLARATIONS

### Data availability

All data used in this project come from published papers as it is described in Materials and Methods.

