## Supplementary Figures for "Causal network perturbations for instance-specific analysis of single cell and disease samples"

Kristina L. Buschur, Maria Chikina, Panayiotis V. Benos
Department of Computational and Systems Biology
University of Pittsburgh School of Medicine
Pittsburgh, PA 15260, USA

### SUPPLEMENTARY FIGURES

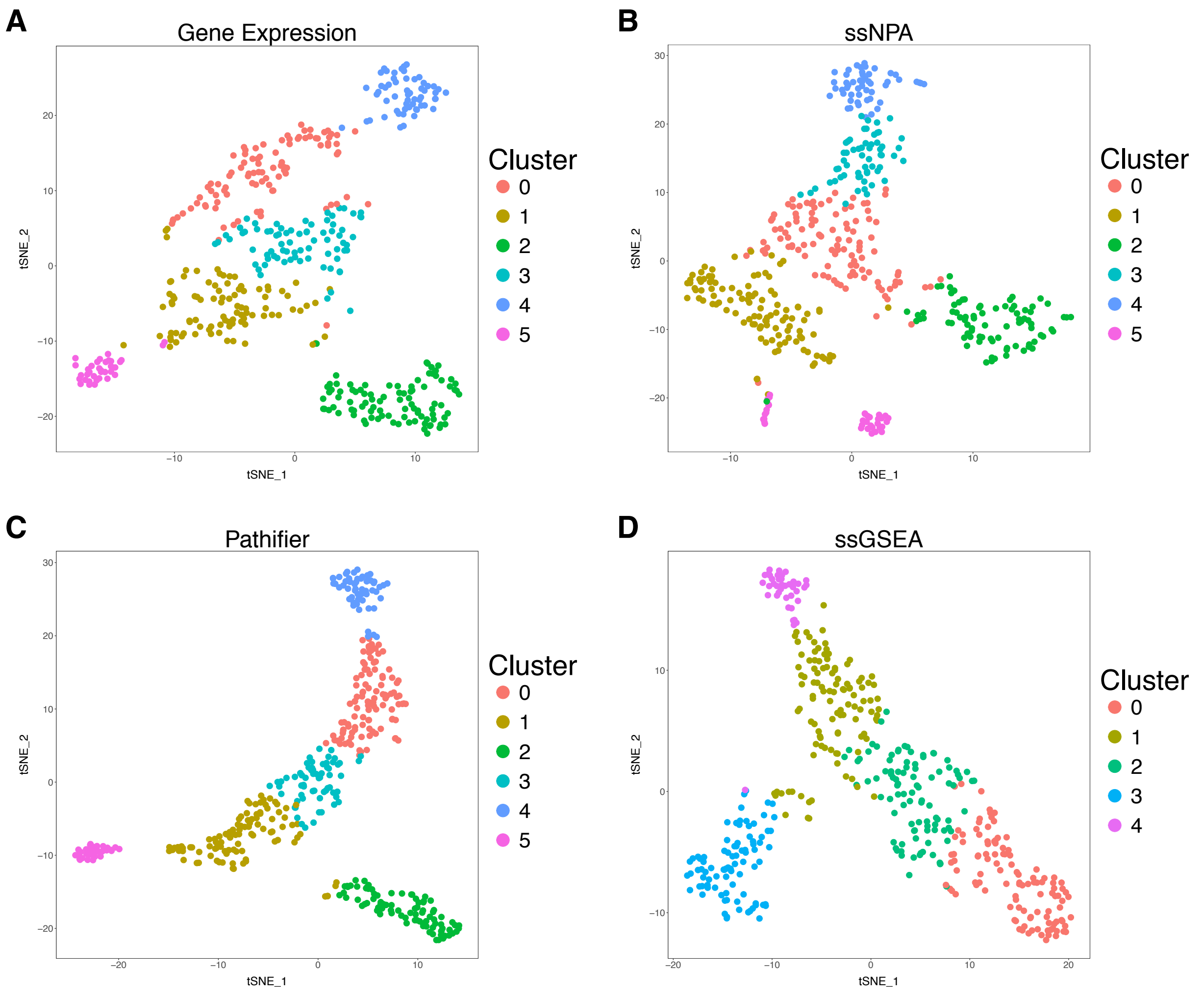


**Supplementary Figure S1.** Cluster assignments with (A) gene expression, (B) ssNPA, (C) Pathifier, and (D) ssGSEA of murine liver cell scRNA-seq samples. ssNPA was used with PD=5 and the E14.5 cells were provided as the reference set to both ssNPA and Pathifier. Clustering for all methods was performed with the first 10 principal components.

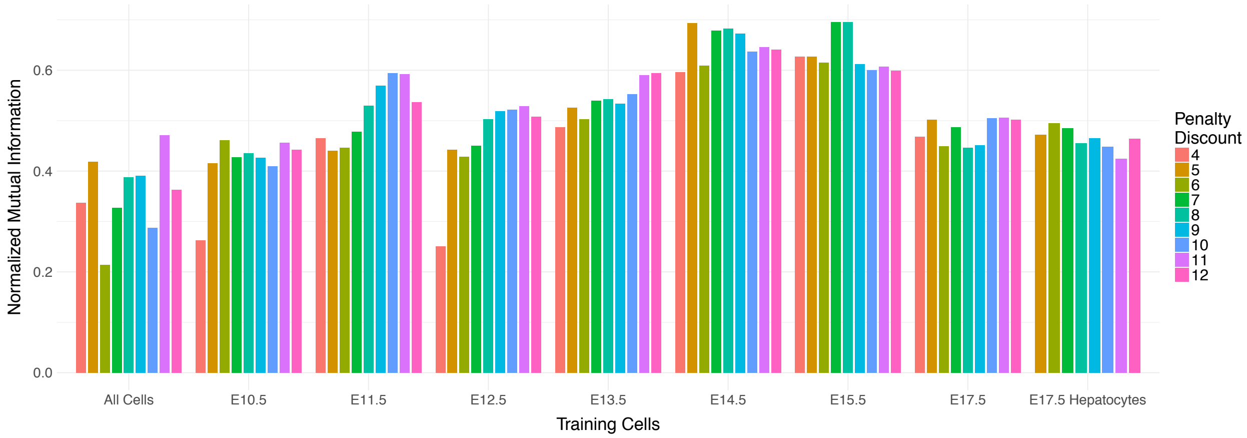


**Supplementary Figure S2.** ssNPA clustering performance assessed by normalized mutual information measures how well murine liver cells are separated by their developmental stage and cell type and is a function of both the penalty discount (PD) parameter chosen for learning the reference causal network with FGES and the cells chosen as the reference set on which to learn the reference network. Clustering for all methods was performed with the first ten principal components. Time point E14.5 with PD=5 were chosen for learning the reference network in all subsequent ssNPA analyses of this dataset.

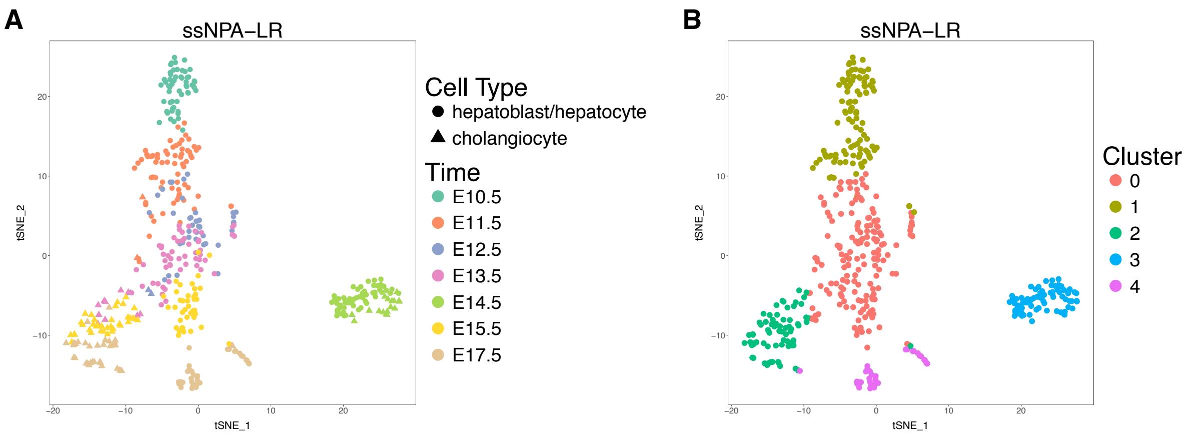


**Supplementary Figure S3.** (A) Developmental stage and cell type separation and (B) cluster assignment with ssNPA-LR on a murine liver cell scRNA-seq dataset. We chose E14.5 as the reference group of cells to facilitate comparison with ssNPA. Sparsity parameters (λ) for the lasso regression models were chosen with 10-fold cross validation, selecting the value of λ corresponding to minimum mean cross-validated error. Clustering was performed with the first ten principal components.
